# Neuronal toxicity and recovery from early bortezomib-induced neuropathy: targeting the blood nerve barrier but not the dorsal root ganglion

**DOI:** 10.1101/2024.05.31.596821

**Authors:** Mariam Sobhy Atalla, Anna-Lena Bettenhausen, Julius M. Verse, Nadine Cebulla, Susanne M. Krug, Reine-Solange Sauer, Mugdha Srivastava, Thorsten Bischler, Jeremy T.C. Chen, K. Martin Kortüm, Robert J. Kittel, Claudia Sommer, Heike L. Rittner

## Abstract

The use of the first in class proteasome inhibitor Bortezomib (BTZ) is highly effective in the treatment of multiple myeloma. However, it’s long-term use is limited by the fact, that most treated patients develop dose limiting painful polyneuropathy. In some of the treated patients, pain resolves after variable timeframes, in others it persists, despite the discontinuation of treatment, with the underlying mechanisms poorly understood. One condition of neural toxicity is the ability to penetrate the blood nerve barrier. Here we present pathways involved in early bortezomib-induced polyneuropathy (BIPN) development and its resolution, in rats and in myeloma patients. One cycle of BTZ elicited transient mechanical hyperalgesia and cold allodynia in rats. Transcriptomic signature and network analysis revealed regulation of circadian, extracellular matrix, and immune genes within the nerve and modest changes in the dorsal root ganglia. Recovery processes resealed the small molecule leakiness of the perineurial barrier, reversed axonal swelling, and normalized small fiber density in the skin. Expression of the microtubule-associated cytoskeletal protein cortactin matched this process in the perineurium. Netrin-1 (Ntn1) as a known barrier sealer was also upregulated in pain resolution in nerve and skin. In patients with painful BIPN skin NTN1 was independent of axonal damage. In summary, our data demonstrate that early BTZ toxicity targets mainly the nerve and indicates that pain resolution could be supported by protective growth factors like Ntn1 for remodeling of the extracellular matrix and neuronal barriers.

**Summary:** Bortezomib leads to dose-limiting painful polyneuropathy. Already in the first cycle, BTZ toxicity weakens the blood nerve barrier which reseals upon upregulation of netrin-1.

## Introduction

Bortezomib (BTZ) is a first-generation proteasome inhibitor described two decades ago for the treatment multiple myeloma and mantle cell lymphoma [39]. Despite its effectiveness, patients experience next to gastrointestinal symptoms and fatigue, in particular hematological and neuronal toxicity, such as thrombocytopenia and peripheral neuropathy [7; 40; 41]. Bortezomib-induced peripheral neuropathy (BIPN) occurs in up to 50% of patients and is characterized by symmetric distal numbness, par- and dysesthesia, neuropathic pain, and weakness. Thrombocytopenia and BIPN often limit the dose and number of cycles administered, resulting in pseudoresistance. In more than 50% of cases, symptoms resolve after dose reduction or therapy discontinuation [41; 44]. Pain resolution is an active process governed by subtypes of immune cells, endogenous opioids, anti-inflammatory cytokines, and specialized pro-resolving mediators [21; 42; 47]. However, pathways and networks regulating this process in the early phase BIPN are unknown.

Recent years have been particularly illuminating on the pathophysiology of BIPN in animal models. Since BTZ does not pass the healthy blood-brain barrier, BIPN is assumed to be driven in the PNS. (Ultra)structural pathology in the nerve has revealed axonal degeneration, mitotoxicity, decreased microtubule dynamics [30; 55] as well as demyelination and Schwann cell degeneration [5; 48]. The immune system is also involved as shown by macrophage infiltration and increased cytokine expression [26; 29]. However, most of the studies concentrated on chronic BTZ treatment (> 4 weeks) and it is unknown which processes occur at the beginning when it is still reversible.

Neuropathy is often initiated by endoneurial oedema and leakage of the blood nerve barrier (BNB).[52] The BNB consists of perineurium, basal lamina, and endoneurial vessels. To secure sealing, cells are connected with adherence, gap, and tight junctions. In perineurial cells, the tight junction proteins ZO-1 (Tjp1) and claudin-1 (Cldn1), are expressed while endoneurial vessels express Cldn5 and Tjp1 [2; 33; 35]. Myelinated nerves are tightened by Cldn19. Intracellular proteins such as cadherin, cortactin (Cttn), and catenin alpha 1 (Ctnna1) link tight and adherence junction proteins to the cytoskeleton.[50] Downregulation of these tight junction proteins results in breakdown of the BNB after traumatic or metabolic (diabetic) nerve injury which goes in parallel with pain and hypersensitivity [2; 8; 27; 36].

Barriers can be re-sealed by several mechanism including netrin-1 (Ntn1) – a chemotropic, secreted, bifunctional guidance cue that directs embryonic PNS development and fosters nerve regeneration [8; 12; 33]. *Ntn1* is expressed in endothelial and perineurial cells and maintains the blood-brain barrier or the BNB by upregulating tight junction proteins.[8; 31] *Ntn1* is also present in the skin [45] presumably nurturing sensory fibers. Furthermore, *Ntn1* in important for the myelin barrier sealing in Schwann cells [8] and vascularization of the developing nerve [49].

Here, we used a single cycle of BTZ treatment to explore early nerve and dorsal root ganglion (DRG) toxicity and its resolution to understand the initial steps in BIPN and its resolution. We further focused on the BNB breakdown and Ntn1 as a barrier sealing and cutaneous neuronal guidance molecule. Finally, we explored similar mechanisms in patients with painful BIPN.

## Methods

### Animal model

All animal care and experimental protocols were approved by the Government of Unterfranken. Male Wistar rats (Janvier Labs) were selected because of the male preponderance in multiple myeloma and BIPN. Rats were injected with BTZ (Selleck Chemicals; 0.2 mg/kg dissolved in 5% dimethyl sulfoxide) or vehicle i.p. twice a week within two weeks (on days 1, 4, 8, and 11) following established treatment regimen in human disease [56]. This schedule imitated one clinical treatment cycle. Thirty minutes after injection, reflexive nociceptive thresholds were determined. Rats were sacrificed and tissue harvested on day 11/12, 18, 25 for transcriptomics, barrier assessment, qPCR, and immunohistochemistry (**Fig. 1a**).

**Figure 1:**
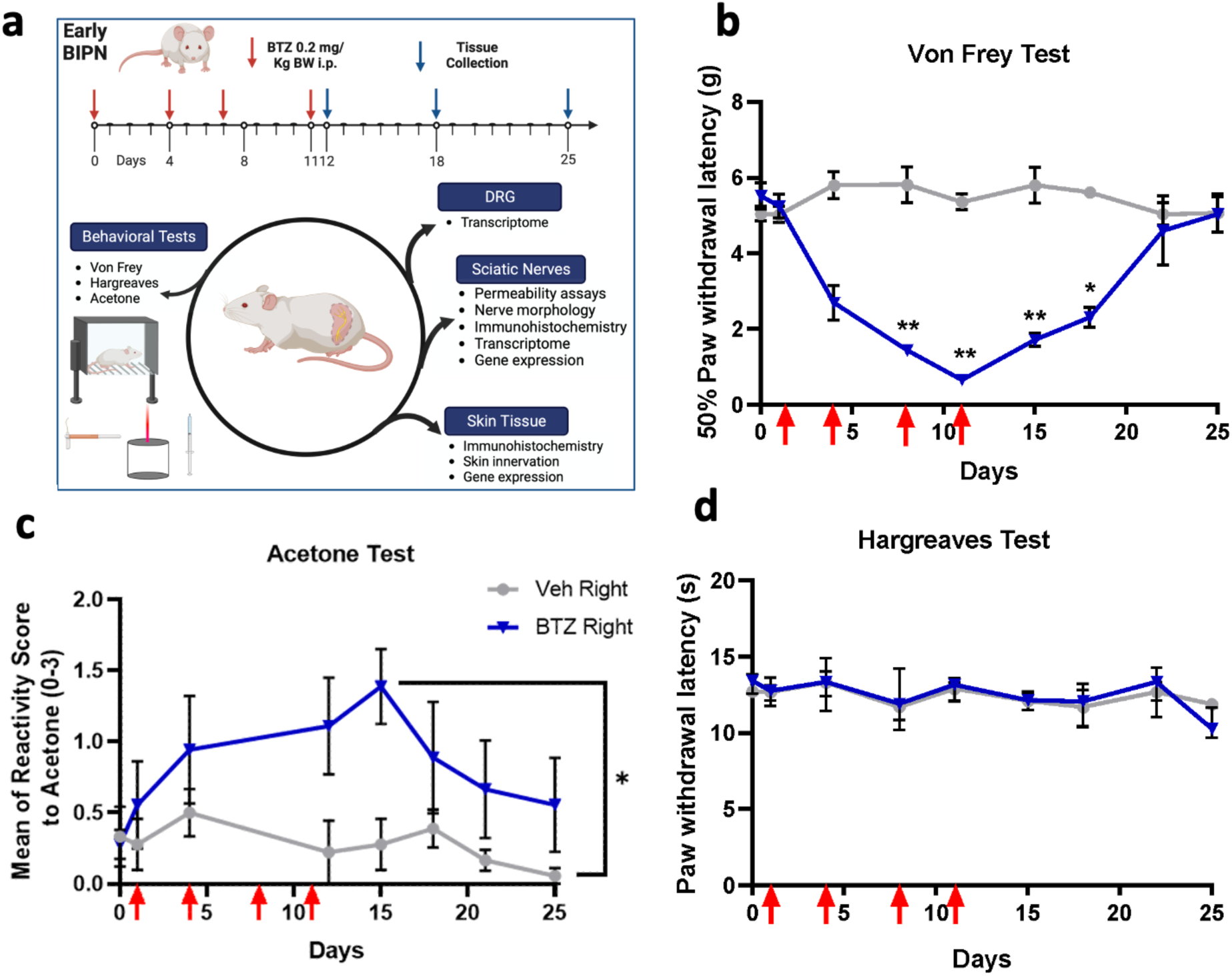
Short-term mechanical hyperalgesia and cold allodynia as well as recovery after on cycle of BTZ. **(a)** Experimental overview: Male Wistar rats were injected four times i.p. with 0.2 mg/kg mg/kg BTZ similar to one cycle in patients and either tested for pain behavior at indicated time points or molecular analysis; control animals were treated with solvent (grey, Veh). **(b)** Mechanical allodynia was measured with von Frey filaments (weight in g), **(c)** cold allodynia by acetone, and **(d)** heat allodynia by Hargreaves test (s). Red arrows depict timepoints of injection. n = 6, all datapoints represent mean ± SEM, RM ANOVA, * p < 0.05, ** p < 0.01.

### Behavioral testing

Animals were randomly assigned to the treatment groups. All experiments were performed by two researchers aware of the treatment group due to the general effects of BTZ [59].

***Mechanical hyperalgesia*** was examined using a series of von Frey filaments to record the withdrawal threshold of the hind paw.[2] We used Dixon’s up and down method to determine the 50% von Frey filament withdrawal threshold using the freely available up-down reader [19].

**Thermal hyperalgesia** was assessed performing the Hargreaves test with a thermal light stimulus. We measured the latency of withdrawal responses with an average from three trials. A recovery period of 10 min was kept and the cut-off latency set to 20 s to avoid any tissue damage [8].

**Cold allodynia** was measured using the cooling effect of acetone: one drop was applied onto the plantar surface of the hind paw [58]. The response was recorded as a numerical score ranging from 0 to 3, where an increase in the numerical value represented a more exaggerated response of paw twitching and licking. An average of 3 consecutive measurements was recorded with 20-30 s intervals between measurements.

### RNA isolation

RNA of the plantar skin and both sciatic nerves nerve was harvested at the indicated time points, immediately shock frozen in liquid nitrogen, and stored then at −80°C. Total RNA was isolated using TRIzol® reagent (Invitrogen, Thermo Fisher Scientific, Darmstadt, Germany) [2] or QIAzol® Lysis reagent (Qiagen, #cat 79306) following vendor’s instructions. RNA concentration and quality were determined using a spectrophotometer (Nandrop-2000, Fisher Scientific). We aimed an optical density quotient (OD 260/280) for follow-up experiments of 1.8 to 2.0.

### Transcriptome analysis

RNA extracted from sciatic nerves and DRGs was used for bulk RNA sequencing. RNA quality was checked using a 2100 Bioanalyzer with the RNA 6000 Nano kit (Agilent Technologies). The RIN for all samples was ≥ 7,7. DNA libraries suitable for sequencing were prepared from 300-500 ng of total RNA with oligo-dT capture beads for poly-A-mRNA enrichment using the TruSeq Stranded mRNA Library Preparation Kit (Illumina) according to manufacturer’s instructions (1/2 volume). After 13-15 cycles of PCR amplification, the size distribution of the barcoded DNA libraries was estimated ∼310 bp by electrophoresis on Agilent DNA 1000 Bioanalyzer microfluidic chips. Sequencing of pooled libraries, spiked with 1% PhiX control library, was performed at 21 to 41 million reads/sample in single-end mode with 100 nt read length on the NextSeq 2000 platform (Illumina). Demultiplexed FASTQ files were generated with bcl2fastq2 v2.20.0.422 or bcl-convert v4.0.3 (Illumina). Sequencing data are available at NCBI GEO (http://www.ncbi.nlm.nih.gov/geo). Raw sequencing reads were quality- and adapter-trimmed via Cutadapt [25] v2.5 using a cutoff Phred score of 20 in NextSeq mode, and reads without any remaining bases were discarded (parameters: --nextseq-trim=20 -m 1 -a AGATCGGAAGAGCACACGTCTGAACTCCAGTCAC). Processed reads were mapped to the rat genome (NCBI RefSeq assembly GCF_015227675.2/mRatBN7.2) using STAR [10] v2.7.2b with default parameters but including transcript annotations from RefSeq annotation version 108 for mRatBN7.2. This annotation was also used to generate read counts on exon level summarized for each gene via featureCounts v1.6.4 from the Subread package [23]. Multi-mapping and multi-overlapping reads were counted strand-specific and reversely stranded with a fractional count for each alignment and overlapping feature (parameters: -s 2 -t exon -M -O --fraction). The count output was utilized to identify differentially expressed genes using DESeq2 [24][1.24.0. Read counts were normalized by DESeq2 and fold-change shrinkage was conducted by setting the parameter betaPrior to TRUE. Differential expression was assumed at adjusted p-value after Benjamini-Hochberg correction (padj) < 0.05 and varying thresholds for log2FoldChange. Volcano plots were generated from DESeq2 results using the Bioconductor package EnhancedVolcano [3]version 1.2.0.

### String analysis

The interaction among differentially expressed genes (with a fold change of at least 1.5 and an adjusted p-val less than 0.05) was investigated using the STRING database. STRING uses text mining, experimental interactions, co-expression, neighborhood, gene-fusion, co-occurrence, and databases information to identify the interactions and calculation of interaction confidence score. Specifically, interactions with a high confidence score of 0.7 or higher were scrutinized, ensuring reliability of the interactions. For better and customized visualization, we imported the resulting networks in Cytoscape. The disconnected genes were removed from the network and only largest connected cluster was considered for visualization. Proteins with shared function in relevant categories were clustered based on literature information. Expression data was mapped on network to visualize the upregulation or downregulation of cluster genes.

### Reverse transcription quantitative PCR

RNA (1 µg, 500 ng for TJPs) was reverse transcribed to cDNA using the high-capacity cDNA reverse transcription Kit (Applied Biosystems, Thermo Fisher Scientific, Darmstadt, Germany) following manufacturer’s instructions. Primers for RT-qPCR were designed by the Primer3 online data bank. For amplification we used the PowerUpTM SYBR® Green MasterMix (Applied Biosystems, Thermo Fisher Scientific, Darmstadt, Germany) and the TaqMan detection system.[2] Next, cDNA was amplified using the 7300 Real-Time PCR System (Applied Biosystems, Thermo Fisher Scientific, Darmstadt, Germany) with the following program for the SYBRGreen method: 95°C for 10 min and 40 cycles at 95°C for 3s and 60°C for 30s and for the TaqMan method: 95°C for 20 s and 45 cycles at 95°C for 1 s and 60 °C for 20 s *Gapdh* (nerve) and *Rlp13a* (skin) were used as a reference gen for quantification. Primer sequences and TaqMan probes are given in **Supplementary Tab. 1 and 2.** Relative abundances to *Gapdh*/*Rlp13a* were calculated by the ΔCt method (threshold cycle value). To compare samples from different groups the comparative cycle threshold (ΔΔCt) method was performed. So finally, mRNA expression was expressed as 2^-ΔΔCt^, standardized to a control condition (vehicle animals) set at 1.

### Semithin sections

For semithin sections, sciatic nerve tissue was fixed in 4% PFA/2.5% glutaraldehyde in 0.1 M NaCaCo solution (pH 7.4). Nerves were postfixed in 2% osmium tetroxide, and embedded in plastic. Semithin sections (0.5 μm) were prepared and stained with azure-methylene blue. Sections were visualized by brightfield microscopy (Biorevo BZ-9000-E, Keyence, Osaka, Japan), blinded and randomized before analysis. Images were processed with Fiji/ImageJ and a semi-automatic quantification tool (gratio.efil.de, Plugin v3.2, setup: 1 px ≙ 0.17761 µm). Nerve fiber density was measured by counting manually myelinated nerve fibers in a defined area of 10.000 µm² (one image/nerve and six animals/group). To evaluate the ratio of axon diameter to total fiber diameter (g-ratio), the plugin algorithm arbitrary marked fibers in the nerve, which were subsequently measured manually. The setup consisted of at least eight measurements per image and four images per animal (n = 6). Measurements continued until at least two axon sizes per quartile were met. In addition, axon diameter as well as myelin area was calculated for fiber distribution studies, assuming nearly round fiber shapes.

### Image analysis for barrier function

Images were blinded and randomized before being analyzed with the open-source program Fiji/ImageJ. Four RGB-8-bit TIF saved images per animal were used for fluorescence measurements. For perineural permeability, the whole endoneurium was manually defined as region of interest (ROI), whereas for Cldn1 fluorescence the whole perineurium was encircled. We calculated the fluorescence intensity (FI) by the subtraction of color intensity (average gray value) minus background signal (mean of color intensity of 3 background areas).[2] The means of four fascicles of each animal were then compared. For the quantification of Cldn1 cell-cell contacts, computational peak analysis was performed subtracting mean gray value*2.5 using the integrated Gaussian blur function (r = 2 pixel).[37] In total, five images per animal were analyzed. Intensity peaks were counted, after drawing a longitudinal line through the center of the perineurium and plotting their intensity maxima (= cell-cell contacts, settings: minimal peak amplitude = 1.5*standard deviation, minimal value of a maximum = 15, minimal peak distance = 1 µm). Subsequently, the number of peaks was normalized to the perineural area and compared between groups.

### Electrophysiological barrier characterization of the perineurial barrier: Dilution and bi-ionic potential measurements

Sciatic nerves were harvested and analyzed in modified Ussing chambers. In brief, sciatic nerves after preparation were placed in cold and gassed transport solution and were kept cooled during transport and preparation. After cleaning and opening the nerve, the inner endoneurial parts were carefully removed and the epiperineurial tissue pieces were mounted into modified Ussing chamber inserts, avoiding including thinned or branched areas.

In the Ussing chambers, voltage and transperineurial resistance (TER; R^t^) were measured. Contribution of the bathing solution to the measured resistance was determined prior to each experiment and subtracted. Ussing chambers and water-jacketed gas lifts were filled with Ringer’s solution (in mM: Na^+^ 140; Cl^−^ 149.8; K^+^ 5.4; Ca^2+^ 1.2; Mg^2+^ 1; HEPES 10; D(+)-glucose 10; D(+)-mannose 10.0; beta-hydroxybutyric acid 0.5; and L-glutamine 2.5 mmol/L; pH 7.4). The solution was equilibrated with 5% CO_2_ and 95% O_2_ at 37 °C.

Sodium and chloride permeabilities were determined from dilution potentials. For this, potential changes were recorded by switching the solution of one hemichamber to a solution containing a reduced concentration of NaCl, but with all other components identical to the standard Ringer’s (osmolality was balanced by mannitol). The resulting dilution potentials were used for further calculations employing the and the Goldman-Hodgkin-Katz equation as described before [20]. Potassium permeability was determined similar in bi-ionic potentials, where the solution in one hemichamber was changed to one in which part of the NaCl was replaced isoosmotically by the KCl.

For the measuring fluorescein fluxes, 5 μL / 5 mL of fluorescein (100 mM) was added to one side of the tissue mounted into Ussing chambers, and samples (300 μL) were collected and replaced 0, 10, 20, 30, and 40 min after addition from the other side. Tracer fluxes were determined from the collected samples by measuring the concentrations with a fluorometer at 520 nm (Tecan Infinite M200, Tecan, Switzerland).

### Immunofluorescence microscopy and IENFD measurement

Harvested nerves were embedded in Tissue-Tek O.C.T compound and snap-frozen in liquid nitrogen. Samples were cryo-sectioned at 10 µm thickness and permeabilized with 0,5% Triton-X100 in PBS solution and blocked for 30 min with 3% bovine serum albumin and 1% goat serum in PBS solution. Antibodies are depicted in **Supplementary Tab. 3.**

Sections were incubated with the primary antibody mouse anti-Cldn1, rabbit anti-Cttn and mouse anti-Ntn1, overnight at 4°C (Supplementary Tab. 3). Washing on the next day was followed by incubation with the secondary antibody goat anti-mouse FITC. Nuclei were stained with DAPI 2 mg/ml in distilled water (Sigma-Aldrich, Ref D9542, 1:3000) for 10 min. Next, slides were rewashed and mounted with Aqua Poly/Mount (Polysciences Inc., Ref 18606). Sections were visualized by immunofluorescence microscopy (Biorevo BZ-9000-E, Keyence, Osaka, Japan) using the same settings for all samples. A higher image resolution was necessary for computational peak analysis, which we obtained by confocal microscopy (Fluoview1000, Olympus).

Skin was fixed in 4% PFA for 1 h and then placed in 10% sucrose overnight followed by Tissue-Tek O.C.T compound. The skin blocks were cryo-sectioned at 40 µm thickness. After 5 min of washing in PBS the skin sections were permeabilized and blocked for 1 h in a 10 % Donkey serum (Sigma Aldrich) and 0.3% Triton-X 100 (Sigma-Aldrich) in PBS solution.

Skin sections were cryo-sectioned at 40 µm thickness and incubated with either rabbit anti-Pgp9.5 and goat anti-collagen IV or mouse anti-Ntn1 antibody overnight at 4°C (**Supplementary Tab. 3**). On the following day the sections were washed and incubated to the corresponding secondary antibodies (**Supplementary Tab. 3**). Next, they were washed again and finally mounted with (Sigma-Aldrich, Ref D9542, 1:3000). Sections were viewed by immunofluorescence microscopy (Axio Imager.M2, Zeiss, Oberkochen, Germany).

### Image analysis for barrier function

Images were blinded and randomized before being analyzed with the open-source program Fiji/ImageJ. Four RGB-8-bit TIF saved images per animal were used for fluorescence measurements. For perineural permeability, the whole endoneurium was manually defined as region of interest (ROI), whereas for Cldn1 fluorescence the whole perineurium was encircled. We calculated the fluorescence intensity (FI) by the subtraction of color intensity (average gray value) minus background signal (mean of color intensity of 3 background areas).[2] The means of four fascicles of each animal were then compared. For the quantification of Cldn1 cell-cell contacts, computational peak analysis was performed subtracting mean gray value*2.5 using the integrated Gaussian blur function (r = 2 pixel).[37] In total, five images per animal were analyzed. Intensity peaks were counted, after drawing a longitudinal line through the center of the perineurium and plotting their intensity maxima (= cell-cell contacts, settings: minimal peak amplitude = 1.5*standard deviation, minimal value of a maximum = 15, minimal peak distance = 1 µm). Subsequently, the number of peaks was normalized to the perineural area and compared between groups.

### Perineurial permeability assessment

To examine the perineurial permeability to small molecules, sciatic nerves were harvested and fixed for 1 h in 4% paraformaldehyde (PFA) followed by 15 min immersion in a 3% sodium fluorescein (NaFlu, molecular weight: 376 Da; Sigma Chemicals) in saline solution 2 ml of Evans blue albumin (EBA, 5% bovine albumin labelled with 1% Evans blue; molecular weight: 68 kDa, Sigma Chemicals) for 1 h.[2] Finally, they were placed in sucrose 10% over night followed by embedding in Tissue-Tek O.C.T compound. After the first third of the nerve was discarded by trimming, 10-µm sections were cut and mounted on microscope cover glasses with permaFlour Mountant (Thermo Scientific).

Ion permeability was examined with electrophysiology of the carefully isolated perineurium in the Ussing chamber.

### Patient cohort

Patients were recruited at the Multiple Myeloma Centre of the Department of Internal Medicine II at the University Hospital Würzburg and are part of a larger monocentric, non-randomized, observational clinical trial (DRKS00025422).[6] All patients underwent a standardized neurological examination and including the Overall Disability Sum Score (ODSS), the modified Toronto Clinical Neuropathy Score (mTCNS) and the Neuropathic Pain Symptom Inventory (NPSI). Nerve conduction studies (NCS) were done on the right sural, median, and tibial nerves following standard procedures and a standardized skin biopsy was taken at the calf and thigh. Only patients with painful BIPN [pain average 2.0 (range 1-6) on the numerical pain scale (0-no pain; 10-worst pain)] were included in the analysis.

### Skin punch biopsy and gene expression

Skin punch biopsy (6 mm; device by Stiefel GmbH, Offenbach, Germany) were taken in local anesthesia from the distal lateral calf according to a published protocol.[51] Skin was stored in 500 µl RNAprotect tissue reagent (QIAGEN GmbH, Hilden, Germany) over night at 4 °C. The next day, the reagent was removed and stored at −80 °C until further processing for gene expression analysis.

The miRNeasy Micro Kit® (QIAGEN GmbH, Hilden, Germany) was used. The samples were thawed on ice and immersed in QIAzol Lysis reagent® (QIAGEN GmbH, Hilden, Germany) using a Micra homogenizer (ART Prozess und Labortechnik, Germany) followed by incubation (room temperature (RT), 5 min). 140 μl chloroform (laboratory production of ultimate elements) was added, shaken vigorously for 15 s and incubated again at RT for 2-3 min followed by centrifugation (12,000 g, 15 min, 4 °C). The upper aqueous phase was transferred to a new collection tube and mixed with 525 µl 100% ethanol (Merck KGAA, Darmstadt, Germany). 700 µl were pipetted into RNeasy MinElute spin column in a 2 ml collection tube, centrifuged (8,000 g for 15 s at RT) and flow-through was discarded. This step was repeated until all of remaining sample was used. 700 μl Buffer RWT was added to the RNeasy MinElute spin column, centrifuged (15 s at ≥ 8,000 g) and the flow-through discarded. 500 μl Buffer RPE were added onto the RNeasy MinElute spin column, centrifuged (15 s at 8,000g) and flow-through was discarded. Next, 500 μl of 80% ethanol (Merck KGAA, Darmstadt, Germany) was added to the RNeasy MinElute spin column, centrifuged (2 min at 8,000 g), flow-through and collection tube was discarded. RNeasy MinElute spin column was placed in a new 2 ml collection tube, centrifuged at full speed for 5 min at RT, flow-through and collection tube was discarded. RNeasy MinElute spin column was placed in a new 1.5 ml collection tube, 14 μl RNase-free water was added directly to the center of the spin column membrane and centrifuged for 1 min at full speed to elute the RNA. After the RNA quality and quantity was assessed with a NanoDrop™ One (Thermo Fisher Scientific, Waltham, MA, USA), the samples were stored at −80 ◦C.

All used cyclers and PCR reagents were purchased from Applied Biosystems (Darmstadt, Germany). For reverse transcription of the obtained mRNA to cDNA, 250 ng of mRNA and TaqMan Reverse Transcription Reagents® were applied in a total volume of 100 µl. The following components were included: 10 µl 10×PCR, buffer, 22 µl MgCl2, 20 µl dNTP, 5µl random hexamer, 2 µl RNase inhibitor, 6.25 µl multiscribe reverse transcriptase. Following PCR cycler conditions were used: annealing (25 °C, 10 min), reverse transcription (37 °C, 60 min), and enzyme inactivation (95 °C, 5 min).

TaqMan Master Mix® and 3.5 μl of sample cDNA were used for real-time qPCR performed in a StepOnePlus™ Cycler. RPL13A (Assay-ID: Hs04194366_g1 VIC-MGB, Applied Biosystems; Darmstadt, Germany), was used as endogenous control. The real-time qPCRs contained 5 µl TaqMan Advanced Master Mix, 0.5 µl TaqMan Assay RPL13A (Assay-ID: Hs04194366_g1 VIC-MGB, Applied Biosystems; Darmstadt, Germany), 0.5 µl of TaqMan Assay Netrin 1 (Assay-ID: Netrin 1 Hs00924151_m1 FAM-MGB, Applied Biosystems; Darmstadt, Germany) and 0.5 µL nuclease free water in a total volume of 6.5 µl. Following PCR cycler conditions were used: incubation (2 min, 50 °C), second incubation (95 °C, 2 min), 40 cycles (3 s, 95 °C and 30 s, 60 °C). We investigated gene expression of Netrin 1 (Assay-ID: Netrin 1 Hs00924151_m1 FAM-MGB, Applied Biosystems; Darmstadt, Germany)

### Statistical analysis

All statistical analyses were done using Graph Pad version 9 software. Given the low number of animals, we used the non-parametric Mann-Whitney-U-test or unpaired t-test for parametric data. For behavioral test analysis, we performed the non-parametric Friedman test RM with Dunn’s *post-hoc* test. Nonlinear regression was proceeded by the Extra sum-of-squares F test, comparing the best-fitting equation models of all datasets. Differences between groups were considered significant if p < 0.05. All data are expressed as mean ± SEM or median ± error.

## Results

### Short-term mechanical hyperalgesia and cold allodynia after one cycle of BTZ

Mechanical hyperalgesia started on day 4, reached a maximum at 11 days (d), and recovered after 25 d (**Fig. 1b**). Pain resolution – reached at 18 d – was defined as an 50% reduction of mechanical hyperalgesia [14]. A similar time curve was observed for cold allodynia, but it was not completely resolved after 25 d (**Fig. 1c**). No heat hyperalgesia was detected (**Fig. 1d**).

### Profiling the sciatic nerve and the DRG in early BIPN

To gain more insight into the local pathophysiology, we profiled the sciatic nerve and the DRG at maximal hyperalgesia at 12 d vs. vehicle control (**Fig. 2a, b**) and 18 d vs. 12d (**Fig. 2c, d**), exemplifying hyperalgesia and pain resolution, respectively. The **three** highly upregulated genes (log2 fold change > -1) belong to circadian transcription factors (*Dpd* and *Nr1d1) and* extracellular matrix proteins *(Cldn11)* (**Supplementary Tab. 4**). Of the five downregulated genes (log2 fold change > 1), three *Thy1*, *Lyve1*, and *Fcgr2a* are associated with the immune system (**Supplementary Tab. 5, Fig. 2d**). *Csf2rb* and *Clec11a* are components of growth factor netwo*rks* also in the hematopoietic system. Thus, BTZ treatment results in local changes in the sciatic nerve but not extensive alterations in the transcriptome with several hundreds of altered genes.

**Figure 2:**
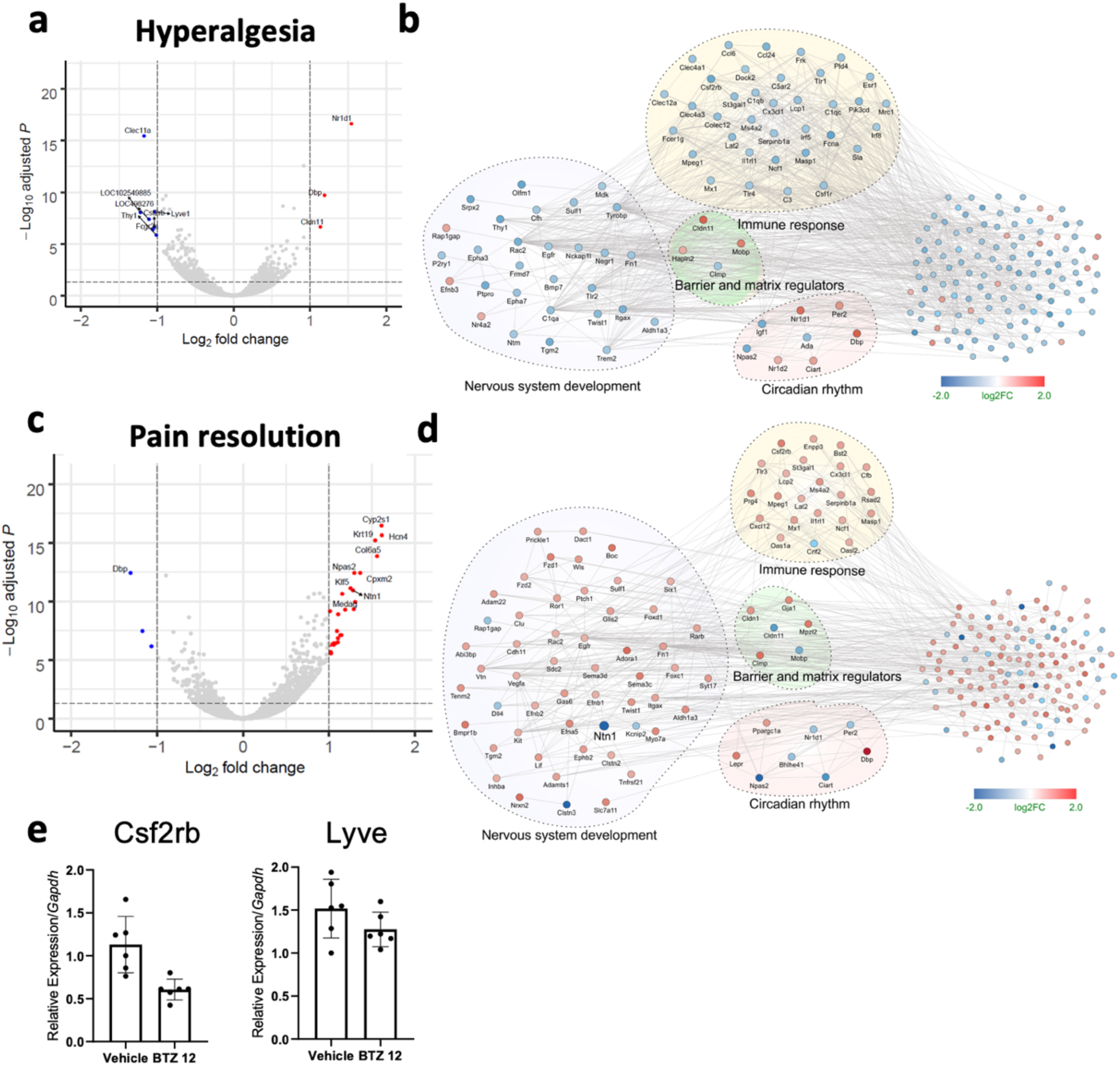
Extracellular matrix regulation, immune systems, and circadian genes as major networks shaping hyperalgesia and pain resolution in BIPN. Sciatic nerves were harvested at the time point of maximum hyperalgesia (12d) and pain resolution (18d) and compared to solvent treated animals (Veh, 12d). (**a, b**) Volcano plots depict up- and downregulated genes at maximum hyperalgesia as well as pain resolution (**Supplementary Tab. 1 and 2**). (**c, d**) Network analysis comparing up- and downregulated genes. n = 3-4/group. (**e**) qPCR validation of Csf2 and Lyve for downregulation at maximum hyperalgesia in independent samples (n = 6, unpaired t-test).

Of the 31 upregulated proteins, almost all (27 genes) were downregulated at pain resolution. Similarly, of the 15 slightly downregulated ones, most were upregulated at 18 d including *Ntn1* and *Hcn4*. So, pain resolution at 18 d was a reversal of previous changes. The spatial and cellular distribution using the sciatic mouse atlas [17] confirmed that BTZ reactivity and remodeling was more prominent in the epiperineurium and the immune system, but Schwann cells were also affected.

We also examined the top differently expressed genes, both up- and down-regulated (long fold change more than 0.7 and les than −0.49) from the samples exhibiting hyperalgesia and pain resolution using dene ontology enrichment analysis (**Supplementary Fig. 1**). Network analysis of the 12 d samples depicted the upregulated circadian networks as well as downregulated immune response regulators (**Fig. 2b**). 18d networks emphasized remodeling with increased cell-cell adhesion and extracellular matrix organization as well as downregulated circadian genes (**Fig. 2d**). The last cluster investigated also included *Ntn1*, a known barrier stabilizer.

In the DRG, we found very few changes (**Supplementary Fig. 2**) – especially no major changes at maximum hypersensitivity. The phase of recovery and pain resolution was also characterized by remodeling and downregulation of extracellular matrix organization.

### Reaction of Schwann cells: reversible axonal swelling but only small changes in myelin

Since some highly regulated genes were *Dpd, Nr1d1, Mpzl2,* and *Mopd* expressed in Schwann cells, we examined the nerve morphology (**Fig. 3**). Nerve fiber density was unchanged with no evidence of axon or myelin degeneration, but the axon diameter increased on average by 20%. Mainly small caliber axons (Ø3.0 ± 0.25 µm) were affected: their overall diameter was reduced by one-fifth, but their axonal diameter increased by 0.5 µm. Both changes normalized in recovery on day 25 (**Fig. 3 d, e**).

**Figure 3:**
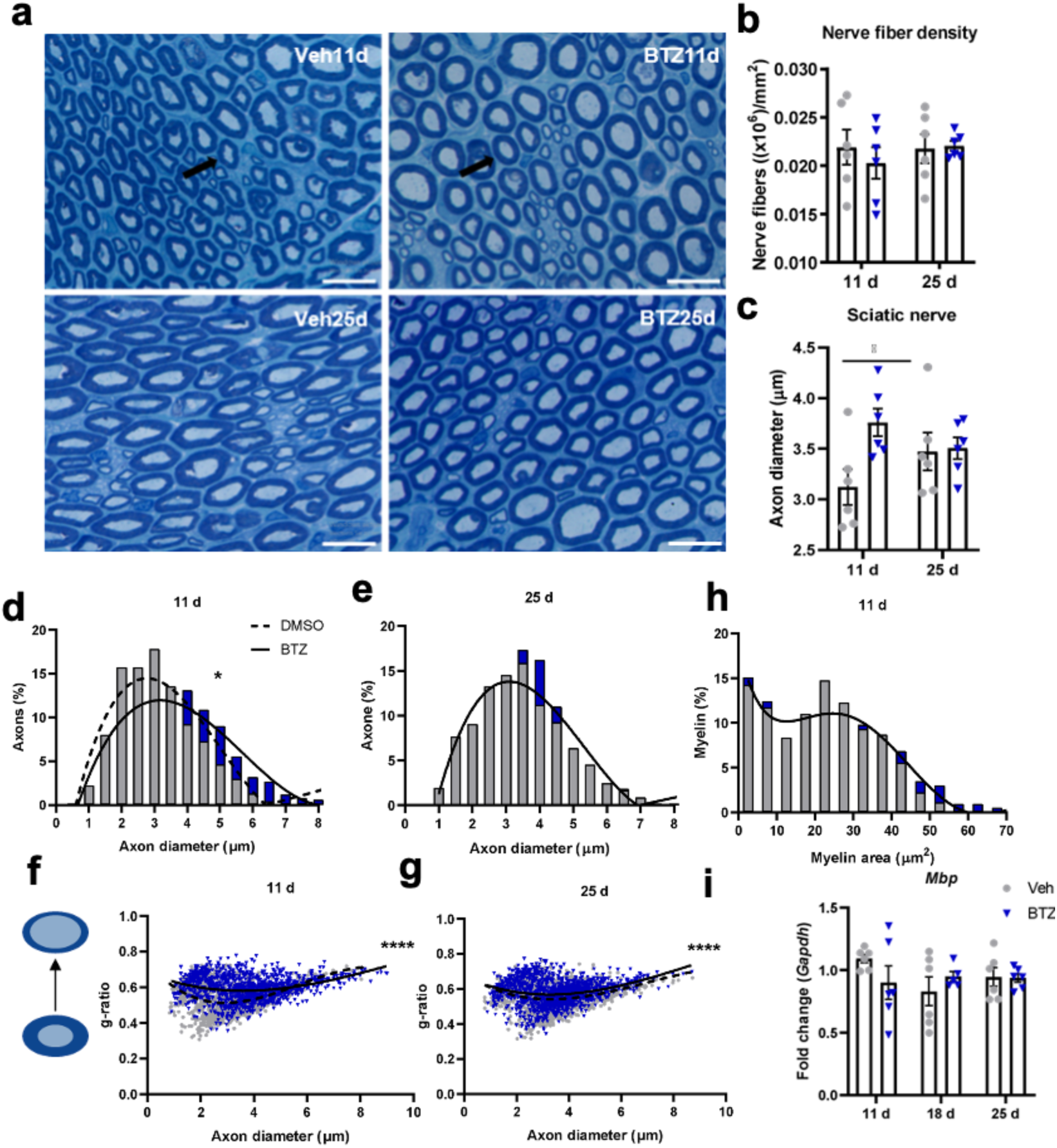
Axonal swelling without myelin alterations in early BIPN. Transient BIPN was induced in male Wistar rats by four injections with BTZ (blue); control animals were treated with solvent (grey, Veh). Sciatic nerves were harvested at maximum hyperalgesia (11 d), pain resolution (18 d) and complete recovery (25 d) and stained with methylene blue. **(a)** Representative plastic sections are depicted. Black arrows point to exemplary axons. **(b, c)** Nerve fibers were quantified in 10.000 µm²/section. Individual axons of the entire nerve section were randomly selected and measured. **(d, e)** The distribution of the axon diameters was analyzed at the time of maximum hyperalgesia and after full recovery. **(f, g)** The distribution and g-ratios: An increase of g-ratio reflects a shift in favor of the axon diameter. Scatter plots show individual fiber datapoints. **(h)** Distribution of myelination at the time of maximum hyperalgesia. **(i)** Mbp was quantified by qPCR. Scale bar: 25 µm. Bar charts represent mean ± SEM, n = 6. * p < 0.05, ****p < 0.0001 always unpaired t-test and F-test for dataset comparisons.

This was reflected in the g-ratio (axon-total fiber ratio) in small caliber axons (**Fig. 3 f, g**). In contrast, the myelin-stabilizing and most abundantly expressed gene myelin basic protein (*Mbp)* was only slightly but not significantly reduced (**Fig. 3 h, i**). In summary, one cycle of BTZ resulted in transient axonal swelling of smaller fibers that resolved after 25 d.

### Increased small molecule perineurial permeability with loss of Cttn and Ntn1

Because of the leakiness of the BNB and specifically the perineurium in traumatic and diabetic polyneuropathy [2; 34; 36; 37], we next wanted to know whether this could also be found in BIPN.

We focused on the epi-perineurium because of changes in the extracellular matrix. After *ex vivo* immersion with fluorescein (330 Da), we observed a 57% higher endoneurial fluorescence signal in BIPN compared to vehicle after 11 d and a complete restoration after 25 d (**Fig. 4 a, b, Supplementary Fig. 3)**. In contrast, the perineurium remained impenetrable for larger molecules like EBA (69 kDa) (**Fig. 4 c, d**). Tight junction proteins like *Cldn1, Cldn5,* and *Cldn19* seal these barriers; *Tjp1* connects them to the cytoskeleton.[35] Expression of all investigated tight junction proteins was unaltered except from *Cldn1* at 11 d (**Fig. 4d-h**). However, immunoreactivity of Cldn1 in the whole perineurium or in cell-cell contacts was unchanged (**Fig. 4i-l**).

**Figure 4:**
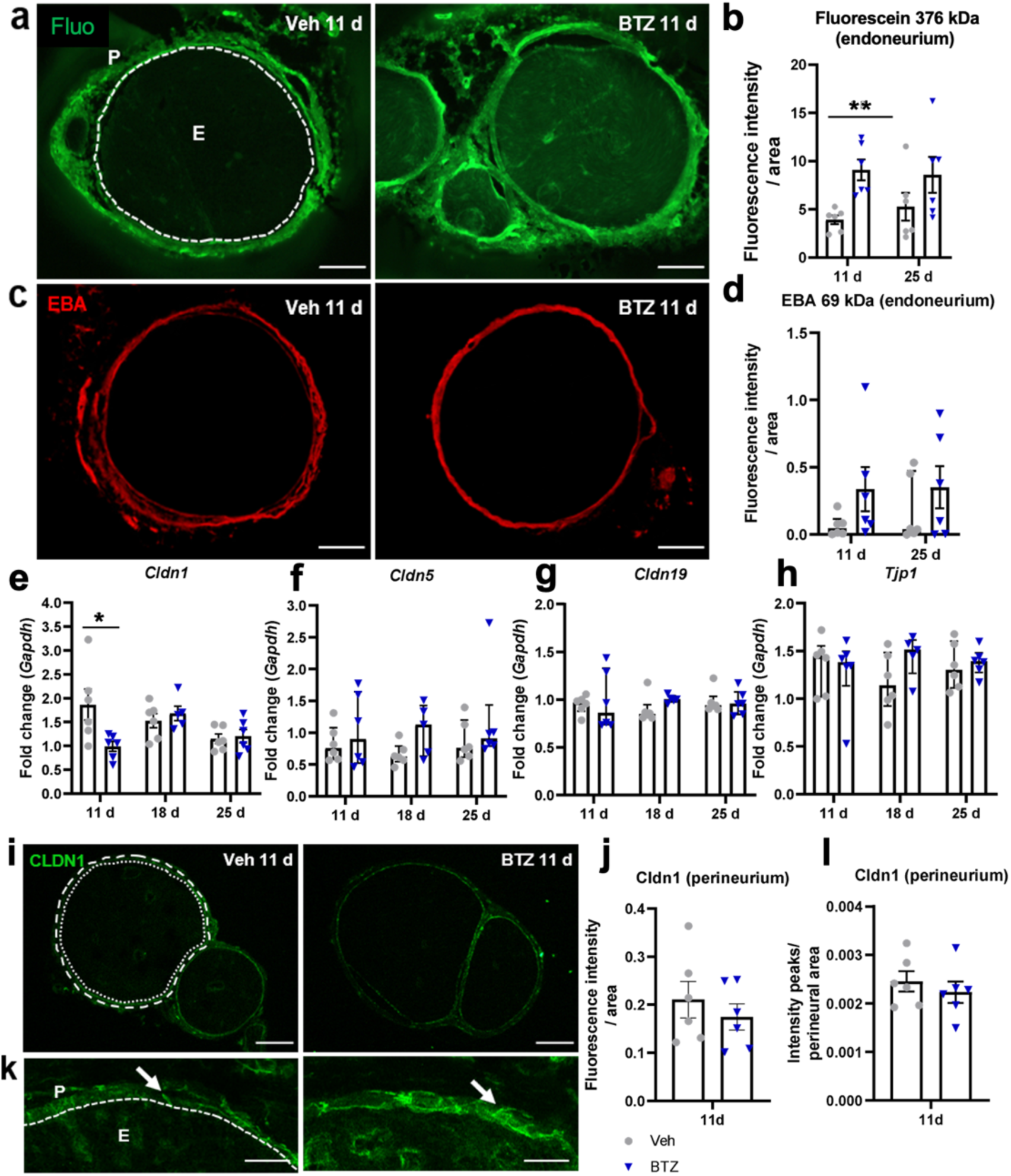
Transient BNB breakdown and resealing for small molecules, but no change in the expression of classical tight junction protein in early BIPN. Male Wistar rats were injected 4-times i.p. with 0,2 mg/kg BTZ (blue datapoints); control animals were treated with solvent (grey, Veh). **(a, b)** Perineurial permeability after ex vivo incubation with fluorescein (Fluo) or **(c, d)** Evans blue albumin (EBA) quantified by fluorescence intensity in the endoneurium (dashed line depicts perineurium). Scale bars = 200 µm. **(e-h)** Relative mRNA expression of different tight junction proteins in the sciatic nerve. **(i, j)** Immunoreactivity for Cldn1 and quantification in the perineurium (fluorescence intensity/perineural area) Dotted line marks the inner border, dashed line the outer border of the perineurium. **(k, l)** Semiquantitative analysis of Cldn1 intensity peaks in the perineurium. Arrows point at Cldn1 in cell-cell contacts. Scale bars = 25 µm. E = endoneurium, P = perineurium. All datapoints represent mean ± SEM, n = 5-6. * p < 0.05 ** p < 0.01, always unpaired t-test except for (d, f-h) Mann-Whitney-U-Test, graphs show median and interquartile range.

Using a novel method to characterize the electrophysiological properties of the epiperineurium ex vivo (**Fig. 5**), we found that permeability for sodium (P_Na_) and chloride (P_Cl_) were unaffected by BTZ, which was also reflected by unchanged transperineurial resistance (TER) that covers all ion permeabilities of the analyzed tissue. However, BTZ had significant effects on the permeability for potassium (P_K_), which was increased after BTZ treatment. After recovery, P_K_ was back to levels as found in the vehicle-treated animals, suggesting a reestablishment of the nerve barrier properties.

**Figure 5:**
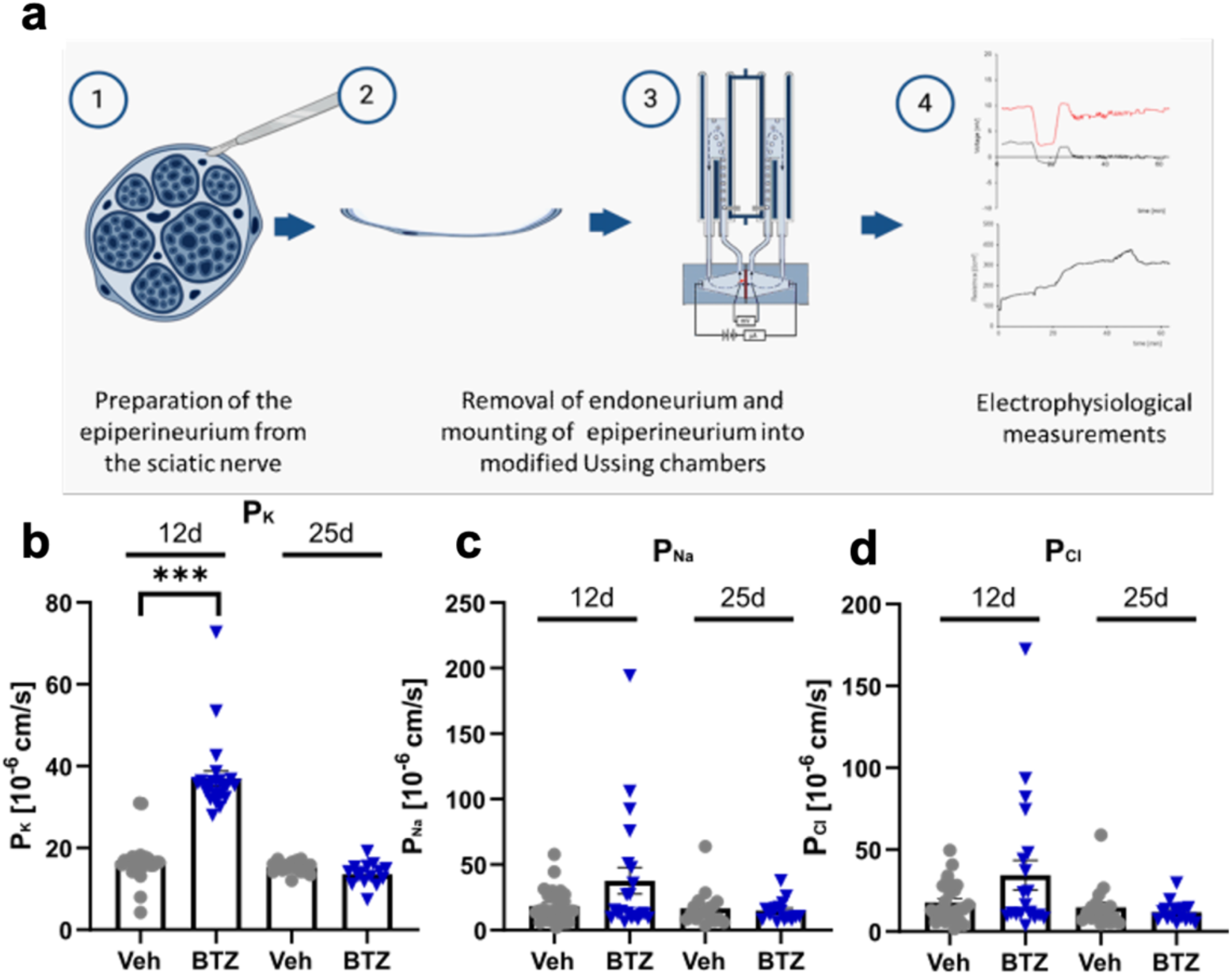
Epiperineurial permeability. BIPN was induced as described before and tissue harvested at indicated time points. **(a)** Workflow to measure perineurial permeability ex vivo. (**b-d**) Ion permeabilities across the epiperineurium were measured at the timepoint of maximum hypersensitivity and at complete resolution. Bars represent mean ± SEM, n = 6-7 (11 d); 3 (25 d) with 8-18 separately measured pieces. ***p < 0.001, unpaired t- test.

Finally, to better understand why the perineurial permeability is increased we searched the literature for alternative pathways. Cttn is an actin binding protein linking cellular membranes to the cytoskeleton to maintain structural integrity also in the BNB.[50] In early BIPN, *Cttn* was significantly reduced only after 11 d (**Fig. 6a**). Immunoreactivity of Cttn confirmed this with an 26% decrease at this timepoint (**Fig. 6b-d**).

**Figure 6:**
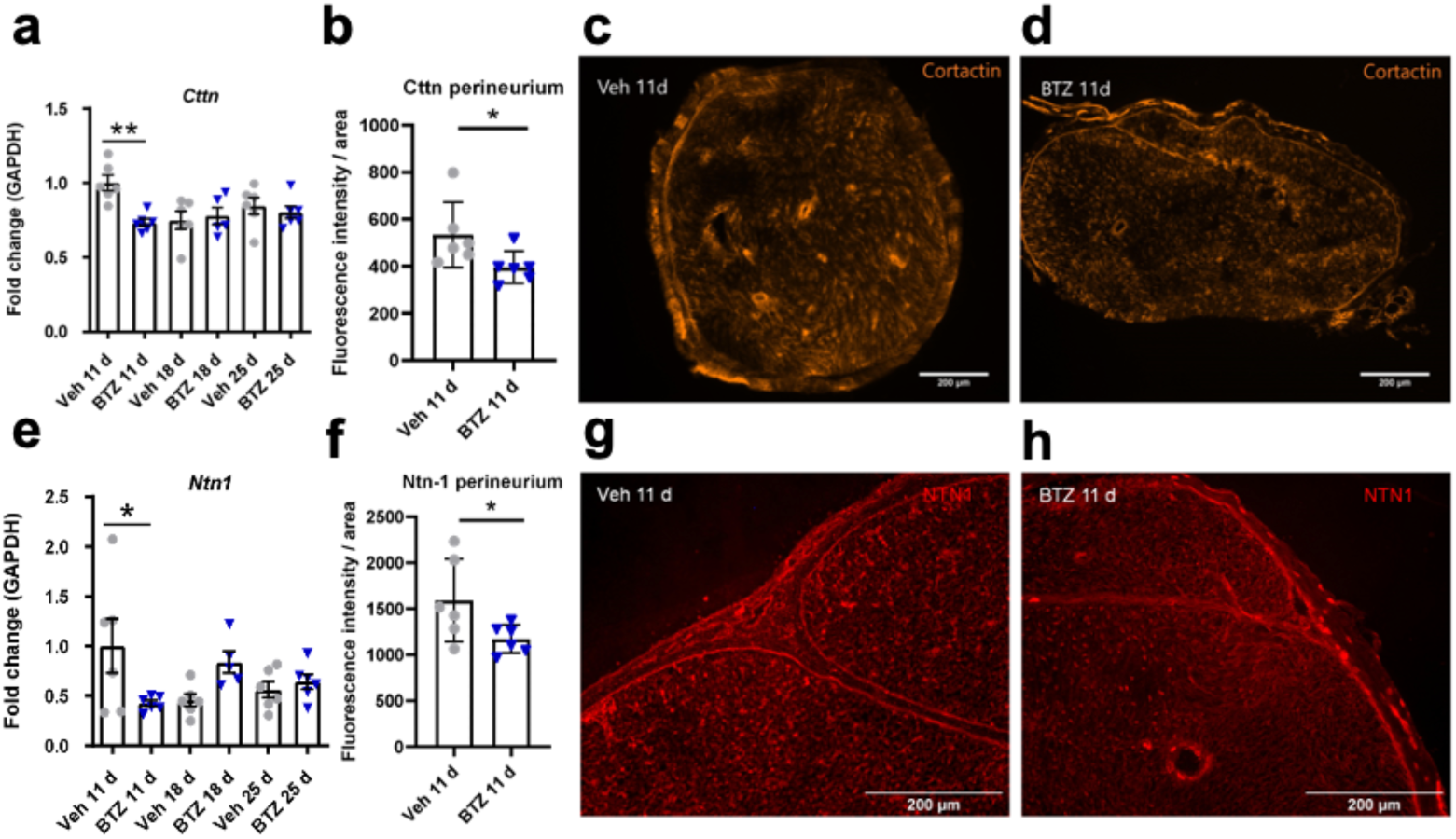
Lack of Cttn and Ntn1 at the timepoint of maximal hyperalgesia and BNB breakdown in the sciatic nerve. BIPN was induced as described before and tissue harvested at indicated time points. **(a, b, e, f)** Relative mRNA expression of Cttn, Ntn1 in the sciatic nerve. **(c, d, g, h)** Immunolabeling for Ctt and, Ntn1 as well as their quantification exclusively in the perineurium. Scale bars = 200 µm. Bars represent mean ± SEM, n = 6. * p < 0.05, ** p < 0.01, two-way ANOVA, Bonferroni post hoc test for multiple comparisons (a, e), or t-test (b, f).

*Ntn1* was upregulated in resolution (**Fig. 2**) and is known to stabilize neuronal barriers.[8] *Ntn1* was downregulated in the sciatic nerve in BIPN by 57% at maximal hyperalgesia (**Fig. 5e**) together with a trend of *Ntn1* increase after 18 d. No change was observed in Ntn1 receptor expression *Unc5b* or *Neo1* (**Supplementary Fig. 4**). Immunoreactivity of Ntn1 in the perineurium was also significantly reduced by 26% in BIPN compared to the vehicle (**Fig. 6e-h**). In summary, hyperalgesia and pain resolution parallel Ntn1 expression.

### Loss of epidermal innervation in parallel with Ntn1 upregulation

To next examine the somatosensory system downstream, we quantified the free endings of sensory neurons: the intraepidermal nerve density (IENFD) in the hind-paws of rats was significantly decreased in hyperalgesia in early BIPN (**Fig. 7a-c**) and recovered at 25d (**Fig. 7d-f**).

**Figure 7:**
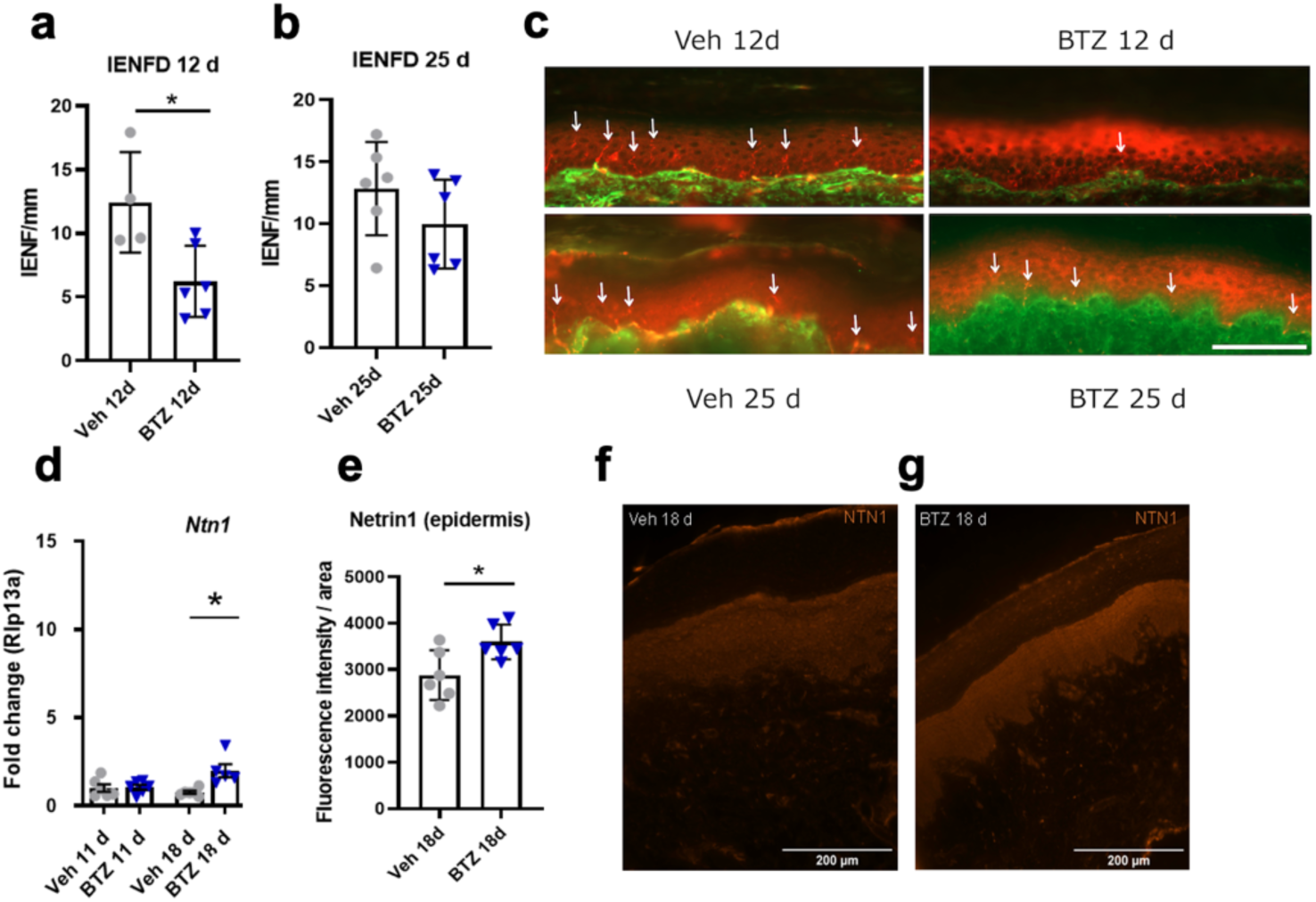
Reduced IENFD at the timepoint of maximal hyperalgesia and upregulation of epidermal Ntn1 in pain resolution. Plantar paw skin tissue was obtained from Wistar rats with BIPN at the time of maximum hyperalgesia (12 d), pain resolution (18 d), and full recovery (25 d). **(a-c)** IENFD was quantified using Pgp9.5 (red) for nerve fiber labelling at the timepoint of maximum hyperalgesia (12 d) and at after full recovery (25 d). Collagen IV (green) immunoreactivity delineates the dermal-epidermal junction. Representative images are shown. **(d, e)** Relative Ntn1 expression in the full planar skin and **(f, g)** immunolabeling for Ntn1 and its quantification in the epidermis in the pain resolution phase at 18 d. Scale bars = 200 µm. Bars represent mean ± SEM, n = 4-6. * p < 0.05 Mann-Whitney-U test.

We hypothesized that this recovering process was correlated with *Ntn1* expression since *NTN1* is also regulated in the skin in patients with neuropathy [45]. *Ntn1* was upregulated in the skin by 160% and immunoreactivity was increased by 25% in the epidermis during pain resolution (**Fig. 6g-j**) while Ntn1 receptor expression was unaltered (**Supplementary Fig. 5**). Ntn1 could be the responsible factor controlling the barrier and somatosensory system.

### Higher Ntn1 in BIPN patients with better sensory nerve function

Finally, we asked whether a shortage of NTN1 is relevant for sensory nerve function in patients with painful BIPN (**Supplementary Tab. 6**). About 1/3 of patients with BIPN reported pain. We selected 16 patients with multiple myeloma and painful BIPN from the ResolvePAIN study [6] with current or past BTZ treatment. Patients with painful BIPN were mainly male, of older age, and suffered from mild to moderate neuropathy (mTCNS) with no muscle weakness (MRC) and mild impairment in daily activities (ODSS). Pain had neuropathic features based on the NPSI. Skin biopsies were taken from the lower calf and NTN1 measured by qPCR. Patients underwent standard electrophysiological analysis including the amplitude of the sensory nerve action potential (SNAP) of the purely sensory sural nerve innervating the lateral ankle and foot to the base of the fifth metatarsal. SNAP reductions are signs of axonal damage. However, when correlating SNAP of the sural nerve and the NTN1 RQ we found no correlation (**Supplementary Fig. 6**). In summary, in a cross-sectional study patient, previous BTZ treatment suffered from neuropathy and a subgroup form pain, but this was independent of NTN1 levels in the skin.

## Discussion

BIPN remains a challenge in the treatment of multiple myeloma because factors governing prevention and recovery as well as pain resolution are yet unestablished. Notably, a single cycle of BTZ is toxic to the nerve – mainly to the epiperineurium – but not to the DRG. In the nerve, the major groups of regulated genes are of circadian, immune, and extracellular protein origin. Pain resolution mostly reverses previous changes and Ntn1 is upregulated. The perineurium is transiently leaky for small molecules especially potassium together with a loss of Cttn. In patients with painful BIPN, epidermal NTN1 was independent of impaired electrophysiological nerve function. Together the preclinical data support the hypothesis that Ntn1 may serve as a possible master regulator for perineurial barrier stability and axonal regeneration in the skin. Its role in patients’ needs to be analyzed in longitudinal studies.

### Loss of Cttn in BIPN accounting for BNB breakdown

A breakdown of the BNB barrier is commonly observed in neuropathies such as chronic constriction injury, spared nerve injury or diabetic polyneuropathy [2; 8; 27; 34; 36; 37]. In traumatic neuropathy models BNB and myelin barrier leakiness are prompted by loss of *Cldn1* (major protein of epi-perineurium), *Cldn5* (main constitute of endoneurial blood vessels), or *Cldn19* (critical for paranodal myelin tightness). In early BIPN, however, only the perineurium was unsealed for small molecules and none of the classical barrier proteins were affected.

Proteins that link tight junction proteins to the cytoskeleton are also important for the structural integrity of barriers, such as the actin binding protein Cttn. A lack of *Cttn* has previously been described in mice with sciatic nerve crush injury. It resulted in a BNB leakage for small molecules [11; 50]. We could confirm this since *Cttn* protein was reduced in the perineurium after one cycle of BTZ. With *Cttn* we identified a molecule that could explain a transient BNB opening.

### Potassium permeability

In the *ex vivo* electrophysiology of the perineurium, BTZ selectively increased the permeability across the perineurium for potassium which recovered after resolution. Potassium is one of the critical factors for proper nerve function and an enhanced permeability leads to ionic imbalance. In damaged nerves, an endoneurial increase of potassium concentration has been reported using ion-sensitive electrodes that was accompanied with spontaneous nerve activity, and that decreased again with ongoing nerve recovery [25] Some BTZ side effects using higher doses suggest a link to impaired potassium concentrations, like hypotension, or bradycardia [15]. In the clinic altered serum potassium concentrations are not common; 0.6-4.1% patients show cardiovascular adverse events probably independent of serum potassium [16]. As inhibitor of the chymotrypsin-like activity of 26S proteasome, BTZ can regulate expression of ion channels including potassium channels and may also lead to surface expression of KATP channels [57]. So, the observed Ionic imbalance could change excitability and foster oedema/swelling.

In our transcriptome analysis *Hcn4* was amongst the top upregulated gene at 18d, the time-point corresponding to resolution. HCN channels are “pacemakers” because they are activated by membrane hyperpolarization, permeable to Na^+^ and K^+^, and constitutively open at voltages near the resting membrane potential. Increased expression and activation of HCN channel isoforms have been implicated in hyperalgesia in oxaliplatin-induced neuropathy [38]. Although *Hcn4* is not involved in neuropathic pain, further studies would need to evaluate its role in resolution [1].

### Reversible axonal swelling

In early BIPN we observed swollen myelinated nerve fibers, but the myelin sheaths were unaffected. This observation is consistent with other studies that found no major destruction of myelin under the treatment with BTZ [18]. Nevertheless, an increase of the cumulative dose leads to Schwann cell dysfunction, myelin aberration, and even the demise of myelinated fibers [26; 55]. Axonal degeneration has been seen in some but not all studies in the early BIPN. Fiber loss is considered unavoidable from a cumulative dose of 3.2 mg/kg. Typical doses of BTZ are 1.3 mg/m² per dose for the d1,4,8,11 scheme (21-day cycle) or 1.6 mg/m² for the weekly d1,8,15,22 in the 28-d cycle. This corresponds to a cumulative dose of 9.36 mg per 21-day-cycle or 11.52 mg per 28-day-cycle. So, clear fiber loss is not likely. Axonal swelling is a hallmark of axonal damage and occurs in mechanical nerve injury and neurodegenerative processes. Impaired axonal transport, mitochondrial dysfunction, microtubule reorganization, and impaired calcium homeostasis trigger axonal swelling [32]. So, axonal swelling in early BIPN could either be due to a decrease in microtubule dynamics with impaired axonal transport [30] and or due to BNB leakage with influx of potassium and other toxic mediators.

### Pain resolution

Several pathways were activated in pain resolution including restoration of extracellular matrix as well as barrier resealing as well as circadian rhythm genes. One important molecule is Ntn1 known as a multifunctional regulator in early neuronal development, Schwann cell proliferation, axon guidance and nerve barrier function [12]. It is downregulated in different types of rodent and human neuropathies [31; 45]. Exogenous netrin restores nociceptive thresholds via perineurial barrier sealing [8]. In early BIPN, lack of *Ntn1* correlates with hyperalgesia and barrier leakiness. The upregulation of *Ntn1* during the recovery phase and the following regeneration of IENFD in the skin underline its role in neuropathic pain resolution in rodent BIPN.

While the circadian rhythm originates in the suprachiasmatic nucleus in the hypothalamus, additional circadian oscillators are found in every cell of the body, including the PNS or the DRG [46]. Altered expression of “clock genes” in sensory tissues contribute to neuropathic pain following peripheral nerve injury, where similar suppression in *Per1*, *Per2*, and *Cry1* expression was observed after partial sciatic nerve ligation in mice [28]. Chemotherapeutic agents such as paclitaxel elicit neuropathy with a clear clinical circadian pattern – suggesting that “clock” genes could regulate the pain response and not simply more attention of patients at night [22]. However, it is quite difficult to differentiate whether the disease, BIPN, or the drug, BTZ, induce such changes. Chemotherapeutic agents, such as BTZ, interfere with the cell cycle and also modify “clock” gene expression, hence affecting the circadian rhythm [43]. Also, melatonin, one of the major hormones regulating the circadian rhythm, inhibits proteasomes: This process interferes with the negative feedback loop of “clock” components such as Cry and Per. So, the proteasome inhibitor BTZ may function in a similar manner [54]. However, very recent evidence rather supports the idea that sensory neurons possess an endogenous clock with regulates the regenerative ability of axons in diurnal oscillation via the core clock protein Bmal1 [9]. Further studies are necessary to dissect this in BIPN.

### Clinical implications

Ntn1 receptors play a key role in tumor progression of several solid tumors: they induce apoptosis when unbound to Ntn1. In fact, dysregulation of these dependence receptors is an important feature of tumorigenesis, because it allows cancer cells to escape apoptosis [4]. Antibodies targeting this interaction (NP137) are currently being evaluated as therapeutic targets [13]. In preclinical and in vitro studies of multiple myeloma, NP137 enhances cytotoxicity of BTZ and dexamethasone and reduces the tumor size. Our data caution the use of these antibodies because they might worsen BIPN or at least hamper their resolution.

In our study we examined early BIPN during only one single cycle of BTZ in rat models that did not develop multiple myeloma. More cycles or the interaction of disease and drug likely trigger further pathways for pain and its resolution. Thus, our study does not completely reflect the clinical situation. Nevertheless, our model resembles BIPN patients suffering from mild neuropathy: BIPN is a predominantly sensory, axonal polyneuropathy with low or absent amplitude of SNAP in the beginning, and reduced amplitude of compound muscle action potentials (CMAP) in more severe cases [53]. NTN1 values tended to correlate with SNAPs of the sural nerve. Although all patients were suffering from neuropathic pain, we only had 1 patient with a severe neuropathy (Severity Grade 3), because patients with severe neuropathy were often not willing to undergo a skin biopsy. Thus, the patients in our cohort were affected by a mild to moderate neuropathy (Severity Grade 1 with pain and 2 with pain), similarly to the animal model.

In summary, early BIPN targets the peripheral nerve and not the DRG with pathophysiological adaptations in ion conductance, extracellular matrix and BNB leakage resolving at the end of the cycle. Ntn1 supports the healing process and pain resolution. Novel pathways like circadian genes and treatment with protective growth factors could be alternative therapeutic targets.

## Acknowledgement

We thank all patients and staff for participation and support of the study. We thank In-Fah M. Lee for excellent technical assistance.

## Data sharing

Data are available on open science data server (Zenodo/GitHub) or with the authors upon reasonable request.

