## Supplemental Figures and Tables for "Neuronal toxicity and recovery from early bortezomib-induced neuropathy: targeting the blood nerve barrier but not the dorsal root ganglion"

### Supplementary tables and figures

| Gene | Company | Number |
|---|---|---|
| *Gapdh* | ThermoFisher Scientific | Rn01462662_g1 |
| *Cldn1* | ThermoFisher Scientific | Rn00581740_m1 |
| *Cldn5* | ThermoFisher Scientific | Rn01753146_s1 |
| *Cldn19* | ThermoFisher Scientific | Rn01416537_m1 |
| *Ocln* | ThermoFisher Scientific | Rn00580064_m1 |
| *Tjp1* | ThermoFisher Scientific | Rn02116071_s1 |
| *Il1b* | ThermoFisher Scientific | Rn00580432_g1 |
| *Il6* | ThermoFisher Scientific | Rn01410330_g1 |
| *Tnfa* (LOC103694) | ThermoFisher Scientific | Rn99999017_m1 |
| *Il10* | ThermoFisher Scientific | Rn01483988_g1 |
| *Mbp* | ThermoFisher Scientific | Rn01399619_m1 |

***Supplementary Table 1 TaqMan-Primers for qPCR***

| Gene | Forward | Reverse |
|---|---|---|
| *Gapdh* | 5′-AGTCTACTGGCGTCTTCAC-3′ | 5′-TCATATTTCTCGTGGTTCAC-3′ |
| *Rlp13a* | 5′-TCTCCGAAAGCGGATGAACAC-3′ | 5′-CAACACCTTGAGGCGTTCCA-3′ |
| *Cttn* | 5′-CTCTCCAAGCACTGCTCACA-3′ | 5′-CGGTCAGCTTGCACTCCATA-3′ |
| *Ntn1* | 5′-CAGGAAGGACTATGCTGTCCA-3′ | 5′-TACGACTTGTGCCCTGCTTG-3′ |
| *Unc5b* | 5′-CGACCCTAAAAGCCGCCCC-3′ | 5′-GGGATCTTGTCGGCAGAGTCC-3′ |
| *Neo1* | 5′-TGTGATGGTGACCAAAGGCA-3 | 5′-GGAGGCTGCCAGTTCACTATT-3′ |

***Supplementary Table 2 Primer sequences for qPCR***

| Antibody name | Target | Vendor | Host organism | Antibody ID |
|---|---|---|---|---|
| Anti-Cldn1 monoclonal antibody (2H10D10) | Cldn1 canine, human, mouse, rat | Thermo Fisher | mouse | AB_2533323 |
| Protein Gene Product 9.5 (PGP9.5, UCHL1) | PGP9.5 human, rat | Zytomed Systems | rabbit | ZYT-516-3344 |
| Anti-type IV collagen-UNLB polyclonal antibody | Collagen IV human, rat | Southern Biotech | goat | AB_2721907 |
| Anti-Cttn polyclonal antibody | Cttn human, mouse, rat | Thermo Fisher | rabbit | AB_2544610 |
| Anti-mouse netrin-1 antibody | Ntn1 mouse, rat | RD-Systems | rat | MAB1109 |
| Anti-mouse IgG FITC polyclonal antibody | Mouse IgG | Sigma-Aldrich | goat | AB_259378 |
| Donkey anti-goat IgG (H+L) cross-adsorbed secondary antibody, Alexa Fluor™ 488 | Goat IgG | Thermo Fischer | donkey | AB_2534102 |
| Donkey anti-rabbit IgG (H+L) highly cross-adsorbed secondary antibody, Alexa Fluor™ 555 | Rabbit IgG | Thermo Fischer | donkey | AB_162543 |
| Goat anti-rat IgG (H+L) cross-adsorbed secondary antibody, Alexa Fluor™ 594 | Rat IgG | Thermo Fischer | goat | AB_10561522 |

***Supplementary Table 3 Antibodies for immunohistochemistry***

| Gene | 12 d | 18 d | Expression in | | | | |
|---|---|---|---|---|---|---|---|
| | | | Schwann cells | epineurial cells | perineurial cells | immune cells | endothelial cells |
| *Nr1d1* | ↑↑ | ↓ | + | ++++ | ++++ | + | ++++ |
| *Dbp* | ↑↑ | ↓↓ | + | ++ | ++ | ++ | +++ |
| *Cldn11* | ↑↑ | ↓↓ | + | + | -++ | - | + |
| *Rph3al* | ↑ | ↓ | - | - | - | +++ | - |
| *Myh7* | ↑ | ↓ | - | + | + | +++ | + |
| *Htra1* | ↑ | ↓ | - | + | + | +++ | - |
| *Mobp* | ↑ | ↓ | + | ++++ | +++ | + | + |
| *Per2* | ↑ | ↓ | + | ++++ | +++ | + | + |
| *Fkbp5* | ↑ | | - | + | + | - | - |
| *Myh7b* | ↑ | ↓ | - | - | - | +++ | - |
| *Htr3a* | ↑ | ↓ | - | ++++ | ++++ | ++ | - |
| *Zbtb16* | ↑ | | - | - | - | - | ++ |
| *Lypd1* | ↑ | | - | ++++ | +++ | - | + |
| *Slc24a2* | ↑ | | - | ++ | +++ | - | +++ |
| *Ryr3* | ↑ | | - | ++++ | ++ | + | + |
| *Ciart* | ↑ | ↓↓ | - | - | +++ | - | - |
| *Galnt15* | ↑ | ↓ | - | - | - | - | - |
| *Htra1* | ↑ | ↓ | - | - | +++ | - | - |
| *Ddn* | ↑ | ↓ | - | - | +++ | - | - |
| *Fxyd7* | ↑ | ↓ | - | +++ | ++++ | - | - |
| *Faah* | ↑ | ↓ | - | +++ | +++ | - | - |
| *Lrg1* | ↑ | ↓ | - | - | + | - | - |
| *Gstp1* | ↑ | ↓ | - | ++ | ++++ | - | + |
| *Bhlhe41* | ↑ | ↓ | - | +++ | +++ | - | ++ |
| *Ttll9* | ↑ | ↓ | - | ++++ | +++ | - | - |
| *Hmgcs2* | ↑ | ↓ | - | - | +++ | - | - |
| *Ca14* | ↑ | ↓ | - | + | +++ | - | + |
| *Rap1gap* | ↑ | ↓ | + | ++++ | ++++ | + | + |
| *Cpt1a* | ↑ | ↓ | - | + | + | - | - |
| *Arhgef3* | ↑ | ↓ | - | - | + | - | - |
| *Pebp1* | ↑ | ↓ | - | - | +++ | - | - |
| *Acaa1a* | | ↓ | - | - | + | - | - |
| *Kcnip2* | | ↓ | + | +++ | +++ | ++ | + |
| *Smoc1* | | ↓ | - | - | +++ | - | - |
| *Abcg2* | | ↓ | - | +++ | ++++ | - | - |
| *Mpped1* | | ↓ | - | ++ | +++ | - | - |
| *Serpinb6b* | | ↓ | - | + | + | - | - |
| *Slc26a8* | | ↓ | - | + | +++ | - | - |
| *Mt3* | | ↓ | - | + | ++++ | - | - |
| *Slc19a3* | | ↓ | - | - | +++ | - | - |
| *Acaa1b* | | ↓ | - | - | - | - | - |
| *Rrad* | | ↓ | - | + | + | - | - |

***Supplementary Table 4 Top genes upregulated at maximum hypersensitivity (12 d) and their recovery during pain resolution (18 d).*** *Location in the sciatic nerve according to the murine sciatic nerve atlas. ↓↓ Log2change <-1, ↑↑ Log2change >1, ↓ Log2change between <-1 and 0, ↑ Log2change between <1 and 0. + slightly, ++ moderately, +++ and ++++ highly, and – not expressed in cells in the sciatic nerve [3].*

| Gene | 12 d | 18 d | Expression in | | | | |
|---|---|---|---|---|---|---|---|
| | | | Schwann cells | epineurial cells | perineurial cells | immune cells | endothelial cells |
| *Clec11a* | ↓↓ | ↑ | - | +++ | +++ | - | - |
| *Fcgr2a* | ↓↓ | ↑ | - | - | - | +++ | - |
| *Csf2rb* | ↓↓ | ↑ | - | - | - | ++ | +++ |
| *Lyve1* | ↓↓ | ↑ | - | + | + | +++ | + |
| *Thy1* | ↓↓ | ↑ | - | ++++ | ++ | + | + |
| *Olfm1* | ↓ | ↑ | + | ++++ | +++ | + | + |
| *Tgm2* | ↓ | ↑ | - | ++ | +++ | - | +++ |
| *Fcna* | ↓ | ↑ | - | - | - | +++ | - |
| *Lum* | ↓ | ↑ | + | ++++ | +++ | + | + |
| *Srpx2* | ↓ | | - | ++++ | +++ | - | + |
| *Itgax* | ↓ | ↑ | - | + | ++++ | - | - |
| *Plxdc1* | ↓ | ↑ | - | - | - | - | - |
| *Adgre1* | ↓ | ↑ | - | - | - | - | - |
| *Qpct* | ↓ | ↑ | - | - | - | - | - |
| *Rac2* | ↓ | ↑ | + | +++ | +++ | + | ++++ |
| *Siglec5* | ↓ | ↑ | - | ++ | ++ | + | + |
| *Npas2* | ↓ | ↑↑ | - | + | + | - | - |
| *Medag* | ↓ | ↑↑ | + | ++++ | +++ | - | - |
| *Cpa3* | ↓ | ↑↑ | - | - | - | - | - |
| *Clmp* | ↓ | ↑↑ | - | ++++ | ++++ | - | - |
| *Hcn4* | | ↑↑ | + | ++ | +++ | + | ++++ |
| *Cyp2s1* | | ↑↑ | + | - | - | - | - |
| *Col6a5* | | ↑↑ | + | + | - | - | + |
| *Krt19* | | ↑↑ | - | +++ | + | - | +++ |
| *Cpxm2* | | ↑↑ | - | - | - | - | - |
| *Clstn3* | | ↑↑ | +++ | +++ | +++ | +++ | +++ |
| *Mettl7b* | | ↑↑ | + | + | ++++ | + | + |
| *Ntn1* | | ↑↑ | + | ++++ | ++++ | + | + |
| *Klf5* | | ↑↑ | + | ++++ | ++++ | + | + |
| *Papln* | | ↑↑ | - | + | + | - | - |
| *Adora1* | | ↑↑ | - | - | ++ | - | - |
| *Nrxn2* | | ↑↑ | - | - | - | - | ++ |
| *Wwc1* | | ↑↑ | - | - | ++ | - | - |
| *Fras1* | | ↑↑ | - | + | ++ | - | - |
| *Slc6a20* | | ↑↑ | - | - | ++ | - | - |
| *Boc* | | ↑↑ | - | +++ | +++ | - | - |
| *Lad1* | | ↑↑ | - | - | ++ | - | - |
| *Mpzl2* | | ↑↑ | - | + | ++++ | - | - |

***Supplementary Table 5 Top genes downregulated at maximum hypersensitivity (12 d) and their recovery during resolution (18 d).*** *Location in the sciatic nerve according to the murine sciatic nerve atlas. ↓↓ Log2change <-1, ↑↑ Log2change >1, ↓ Log2change* between <*-1* and 0, ↑ *Log2change* between <*1* and 0. *+ slightly, ++ moderately, +++ and +++ highly, and – not expressed in cells in the sciatic nerve [3].*

| **Patients with painful BIPN** | N = 16 |
|---|---|
| Sex (Male/Female) | 13/3 |
| Severity Grade 1 with pain<br>Severity Grade 2 with pain<br>Severity Grade 3 with pain | 6<br>9<br>1 |
| Age in years: Median (Range) | 67.0 (50-82) |
| ODSS: Median (Range) | 3.5 (0-6) |
| MRC: Median (Range) | 118 (112-120) |
| mTCNS sum score: Median (Range) | 18.5 (9-27) |
| NPSI sum score: Median (Range) | 21 (3-91) |
| Disease duration in months: Median (Range) | 45 (2-205) |
| Cumulative dose BTZ ($mg/m^2$): Median (Range) | 22.2 (3.9-98.8) |
| Time since last application (months): Median (Range) | 2.5 (0-77) |

***Supplementary Table 6 Patient cohort***. *Severity Grade of neuropathy was graded based on [2]; ODSS – Overall Disability Sum Score (range 0-12 – severest impairment). MRC – Medical research council for muscle strength (range 0-5; 5-maximimun strength; sum score 0-120). mTCNS – modified Toronto Clinical Neuropathy (range 0-33) [1]; NPSI – Neuropathic Pain Symptom Inventory (0-100; 100 maximal neuropathic symptoms).*

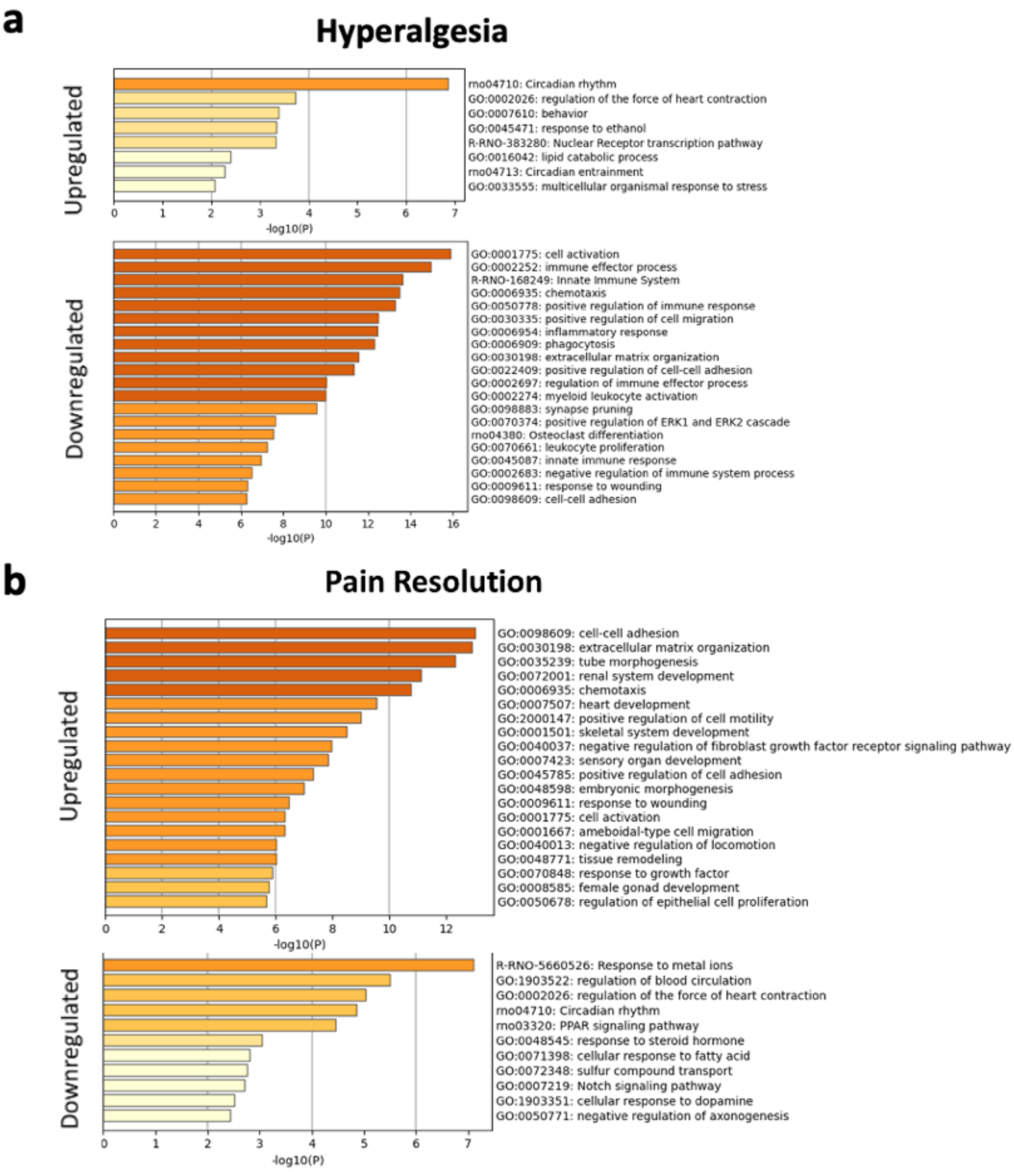


***Supplementary Figure 1: Pathway analysis in the sciatic nerve in BIPN. (a, b)*** *GO Term analysis of up- and downregulated genes comparing. n = 3/group.*

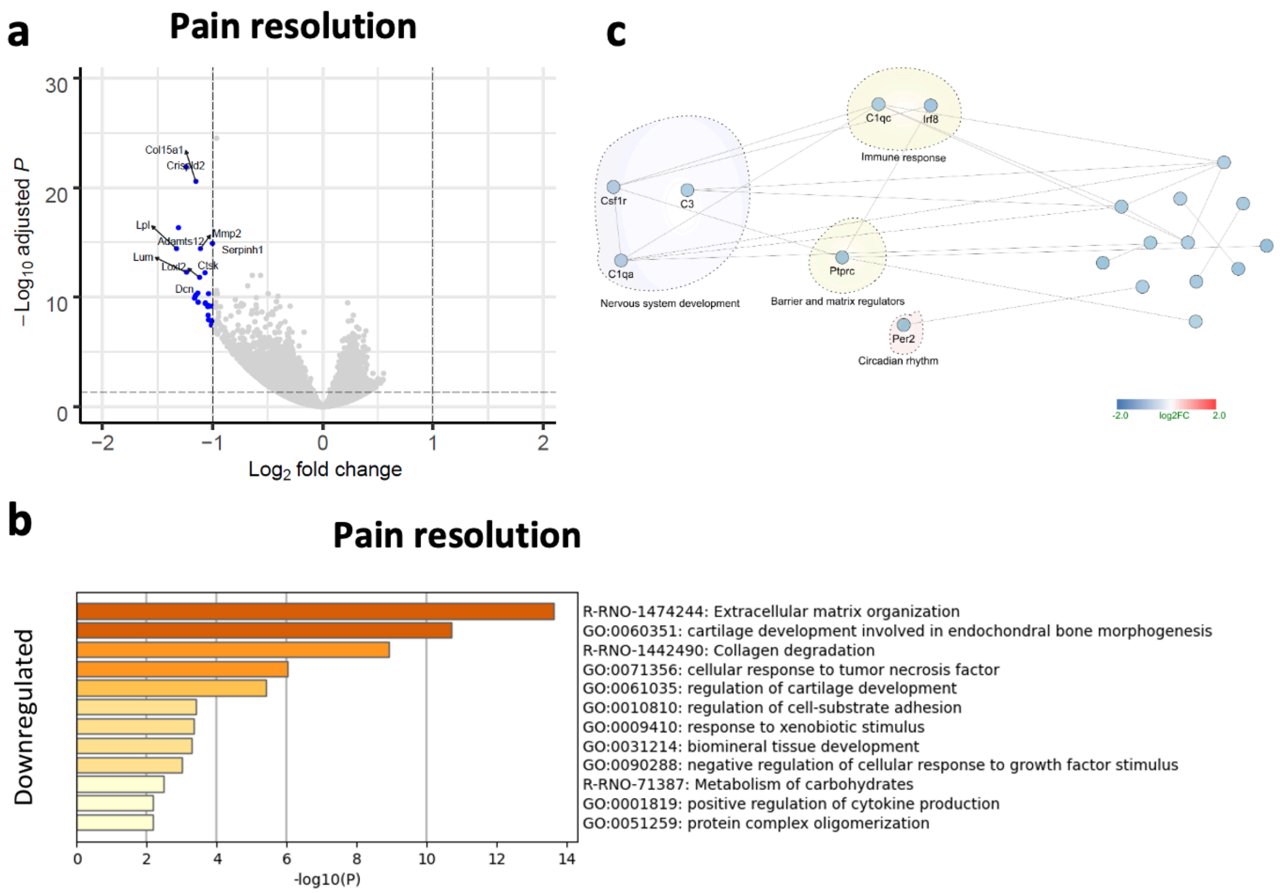


***Supplementary Figure 2: Less network changes in DRGs in pain resolution.*** *DRGs were harvested at the time point of maximum hyperalgesia (BTZ_12d) and pain resolution (BTZ_18d). No changes were found at maximum hyperalgesia compared to vehicle control. (**a**) Volcano plots depict up- and downregulated genes for pain resolution comparing BTZ_18d with BTZ_12d. (**b, c**) Pathway and network analysis of downregulated genes. n = 6.*

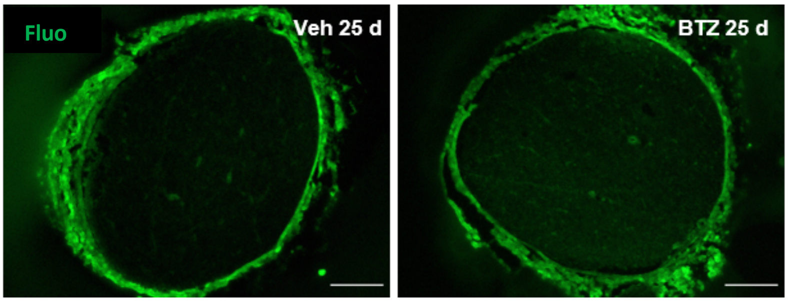


***Supplementary Figure 3: Restored blood-nerve-barrier after complete recovery of BIPN.*** *Perineurial permeability after ex vivo incubation with fluorescein (Fluo, 330 Da). Scale bar = 200 µm.*

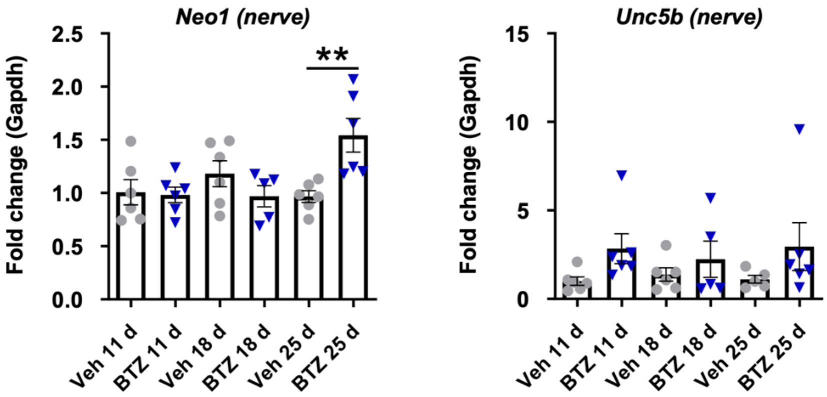

***Supplementary Figure 4: Relative mRNA expression of Ntn1 receptors in the nerve**. All datapoints represent mean ± SEM, n = 5-6. ** p < 0.01, two-way ANOVA, Bonferroni post hoc test for multiple comparisons.*

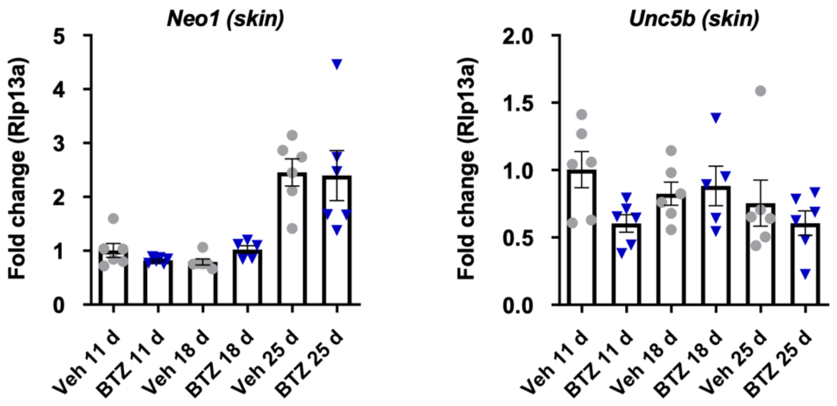


***Supplementary Figure 5: Relative mRNA expression of Ntn1 receptors in the skin**. All datapoints represent mean ± SEM, n = 5-6. p > 0.05, two-way ANOVA, Bonferroni post hoc test for multiple comparisons.*

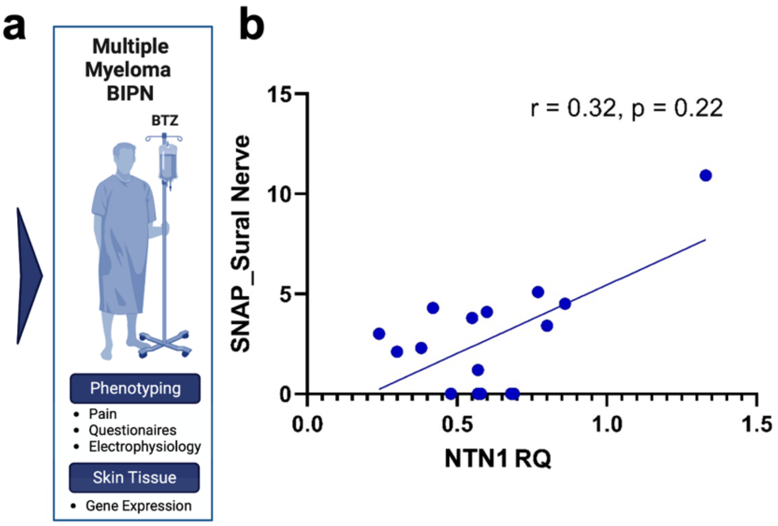


***Supplementary Figure 6: NTN1 expression and sural nerve action potentials.** (**a**) Summary of analyses in patients (**b**) NTN1 was quantified in relation to a calibrator in skin samples from lower calf in patients with painful BIPN. NTN1 RQ values were correlated with the size of the action potential of the sural nerve. Spearman correlation. n = 16.*
